# Indirect restoration of ecological interactions: reintroduction of a dung-beetle associated primate increases the recruitment of large seeds

**DOI:** 10.1101/2021.01.20.427460

**Authors:** Anna R. Landim, Fernando A. S. Fernandez, Alexandra S. Pires

## Abstract

The biased loss of large and medium frugivores alters seed dispersal and plant regeneration. Species reintroductions have been proposed as a strategy to reverse the consequences of species loss. However, the effects of reintroductions on ecological processes are seldom accessed, which hinders the comprehension of reintroductions’ potential to reestablish functioning ecosystems. In this study, we investigate the effect of howler monkey (*Alouatta guariba*) reintroduction on the plant regeneration of Tijuca National Park (TNP), a defaunated Atlantic Forest fragment. Howlers are folivore-frugivore primates, whose large clumped defecations attract dung beetles, which provide secondary dispersal by burying seeds present in the howlers’ feces. Thus, we expect that the fate of seeds dispersed by howlers will differ from those dispersed by other frugivores present in the Park. We followed the fate of seeds between 3 and 14mm in diameter in three steps of the seed dispersal loop, each one consisting of a different experiment. First, we estimated secondary seed dispersal and burial depth probabilities according to the frugivores’ defecation pattern; then, predation probability in different burial depths and defecation patterns; and, finally, recruitment probability in different burial depths. Considering the final result of the three experiments, the howlers’ reintroduction affected positively the regeneration of large seeds. The fate of 3mm seeds was little affected because they were seldom preyed upon at shallower depths anyway and could not recruit when deeply buried. On the other hand, seeds larger than 3mm reached the seedling stage more frequently when dispersed by howlers than when dispersed by other animals present in the Park. Thus, howler monkey reintroduction in defaunated areas, in addition to smaller frugivores, whose defecation patterns are less attractive for dung beetles, improves the regeneration of large seeds. We hope that this study will stimulate new howler reintroductions in defaunated areas.

## 1 Introduction

Human activities have been affecting biodiversity for over 50.000 years (Koch and Barnosky 2006, Araujo et al. 2017). Larger vertebrates are particularly vulnerable due to hunting, habitat degradation and fragmentation (Peres 2002, Peres and Palacios 2007, Pardini et al. 2010, Ripple et al. 2019). This bias is of major concern, since body size partially determines the contribution of species to some ecosystem processes, such as seed dispersal (Vidal et al. 2013, Schleuning et al. 2015). Larger frugivores interact with larger seeds and disperse them through longer distances (McConkey and Brockelman 2011, Dehling et al. 2016). Thus, downsized frugivore communities cannot fully substitute functionally the extinct large and medium-sized animals. The reduction of seed dispersal affects plant recruitment and population growth (Traveset and Riera 2005), modifying plant community composition (Terborgh et al. 2008).

In the present biodiversity crisis, conserving what is left is not enough. The reintroduction of locally extinct species to restore interactions and their associated functions has been proposed to reverse the consequences of species loss (e.g. refaunation, (Oliveira-Santos and Fernandez 2010) and trophic rewilding (Svenning et al. 2016)). Just as the extinction of species leads to losing interactions, species reintroductions can rewire these interactions and reestablish ecological processes (Genes et al. 2019, Mittelman et al. 2020). Nevertheless, most reintroductions are focused on species conservation and the effect of reintroductions on ecological and ecosystem functions is seldom evaluated (Polak and Saltz 2011).

A possible way to evaluate the effect of reintroductions on ecological processes is to estimate the number of restored interactions (Genes et al. 2017). However, two animals can interact with the same seed species but lead to different seed fates (Culot et al. 2017, Lugon et al. 2017). Another way to evaluate the contribution of a newly reintroduced frugivore is to compare the fate of the seeds they disperse with those dispersed by the previously present frugivores. Nevertheless, accompanying the seed dispersal loop is challenging. This challenge can explain the lack of studies that follow many steps of seed fate (Wang and Smith 2002) or that adopt a community perspective and include more than one taxon (Andresen et al. 2018).

Primates are some of the last large mammals remaining in the tropics and among the most threatened groups (Estrada et al. 2017). Although they constitute a large proportion of the frugivore biomass in tropical forests (Eisenberg and Thorington 1973), only recently the number of primates’ seed dispersal studies has increased (Andresen et al. 2018). In the Atlantic Forest, primates interact with all seed sizes (Bufalo et al. 2016), have a generally positive effect on seed germination (Fuzessy et al. 2016) and disperse seeds to long distances (Fuzessy et al. 2017). Nevertheless, there are still many gaps in the knowledge of their role on plant regeneration (Chapman and Dunham 2018).

Howler monkeys (*Alouatta* spp.) are folivore-frugivore medium sized primates. They swallow up most of the seeds from the fruits they eat and affect positively seed germination (Arroyo-Rodríguez et al. 2015). Howlers usually defecate simultaneously, which generate large clumped defecations and high seed densities (Julliot 1996). This pattern attracts rodents and other seed predators, but also dung beetles (Scarabaeidae: Scarabaeinae) (Andresen 2002a). Dung beetles feed on feces and bury seeds while building their nests, a process known as secondary seed dispersal (Andresen and Feer 2005). Secondary seed dispersal alters seed survival and recruitment probability (Shepherd and Chapman 1998). While seed burial decreases predation by rodents, it decreases seedling recruitment, often to very low rates when below 3cm in depth (Andresen and Feer 2005). Thus, when assessing the role of seed dispersers, it is essential to take into account the secondary seed dispersal probability associated with each primary disperser (Culot et al. 2018).

In this study, we investigated the effect of the reintroduction of red howler monkeys (*Alouatta guariba*) on plant regeneration in a defaunated Atlantic Forest fragment. We compared the fate of seeds dispersed by howlers with other frugivores already present in the fragment through three experiments. First, we estimated the probability of secondary seed dispersal according to the frugivore’s defecation pattern; then, we estimated the predation probability for different defecation patterns and burial depths; and finally, the recruitment probability in different burial depths. Each step consisted of a different experiment. We hypothesized that the effect of howlers on plant regeneration, through their clumped defecation, would differ from the effects of the other frugivores already present in the fragment.

## 2 Methods

### Study site

This study was carried out from September 2019 to January 2021 at Tijuca National Park (TNP, 3.953 ha). Mean temperature varies between 18 and 26°C, with rainy summers, surpassing 1.200mm yearly (ICMBIO 2008). TNP is situated within the city of Rio de Janeiro and its history is intertwined with the city’s. The area that now corresponds to the Park used to be explored for coffee farming and coal production until the middle of the 19th century, when it began to be reforested (Padua 2002). Despite some exotic species introduced during reforestation, the vegetation is mainly composed of native Atlantic Forest species (Freitas et al. 2006). Since the TNP is surrounded by a metropolitan matrix, the missing fauna has been unable to recolonize. Although there are still seed dispersers present (e.g. the lowland paca (*Agouti paca*) and the brazilian squirrel (*Sciurus aestuans*)), the Park lacks medium and large frugivores. Recently, three vertebrate species have been reintroduced to TNP, starting with agoutis (*Dasyprocta leporina*) in 2010 (Cid et al. 2014), followed by howler monkeys (*Alouatta guariba*) in 2015 (Genes et al. 2019) and, more recently, in 2019, yellow-footed tortoises (*Chelonoidis denticulatus*).

### The experiment

We compared the seed fate of seeds dispersed by howler monkey’s and the mammal and avian species at TNP whose fruit diet most overlapped with that of howlers’: capuchin monkey (*Sapajus nigritus*) and rusty-margined guan (*Penelope superciliaris*). Diet overlap was based on data from the Atlantic Frugivory data set (Bello et al. 2017), the most complete set of frugivory interactions from the Atlantic Forest so far. The selection of seed species was based on the howler’s credit of ecological interactions in the TNP (Genes et al. 2017), that is, the list of TNP’s plant species with which the howler has the potential to interact. We chose species of plants whose genera (or species) were also part of the capuchin monkey and guan diet. Additionally, we only selected tree species of recalcitrant seeds that were dispersed by zoochory.

Three experiments were set to follow the seed path from seed deposition by the primary disperser (howlers, capuchins or guans) to its recruitment into seedling. The seed fate and predation experiments were both set in TNP, in the same plots, 48hr apart; whereas the seedling recruitment experiment was set indoors. In order to understand if the howlers’ effect on seed fate and seedling recruitment varies with seed size, all experiments were assembled for seeds with four diameters: 3mm (*Psidium cattleianum*, Myrtaceae), 7mm (*Cordia superba*, Boriginaceae), 10mm (*Nectandra membranacea*, Lauraceae) and 14mm (*Virola bicuhyba*, Myristicaceae). All seeds used in the experiments were donated from partner greenhouses.

#### Seed burial and predation

We set up 120 stations in TNP, 50m apart. Each station contained a combination of one seed/bead size and one primary disperser treatment. Primary disperser treatments were set according to the disperser feces deposition pattern. Howler’s feces were placed in plots of 2m^2^ in 10 piles containing 10gr each. Capuchin’s feces were set in 100m transects each, with 8gr of feces (Mikich et al. 2015) every 10m. Guans’ feces were also set in 100m transects, but with 7gr in each pile (Mikich 2002). Stations were replicated 10 times for each seed/bead size and primary disperser combination (but we only replicated the *V. bicuhyba*’s predation experiment 5 times due to limitations in seed availability). In total, we used 1.500, 900, 600 and 300 beads of 3, 7, 10 and 14mm of diameter, respectively, 1500 seeds of *P. cattleianum, C. superba* and *N. membranace*a, and 450 seeds of*V. bicuhyba*.

Seed burial experiments occurred from September to November 2019. We collected howlers’, capuchins’ and guans’ feces from populations at Rio de Janeiro’s Zoo in the mornings. The feces were used on the same day in the experiments at TNP. Since in this first step we only wanted to analyse the probability of secondary seed dispersal and depth, we used plastic beads with the seeds’ diameters (3, 7, 10 and 14mm). We varied the number of beads in the piles of feces according to their sizes: smaller beads were more abundant than bigger ones. In each pile one bead was tied with a 20cm nylon thread with a colored tape in the tip. We returned to the experiments 48hr after they were set, to check if the beads had been buried or not. When positive, we would also measure the burial depth.

Seed predation experiments were arranged following the data collection from bead burial. Seeds were buried 1, 3, 5 and 10cm deep, 20cm from each other, at the same location where the dispersers’ feces had been set. We also included one unburied seed to estimate the probability of predation with no secondary seed dispersal. Seeds were cleaned of their pulp and glued (Tek-bond 793) to a 20cm nylon thread with a colored tape in the tip, and left to dry for over a week. We checked the experiments every month for 6 months. Seeds were considered as preyed upon if they were not found in a 1m radius, if we found the nylon thread but the seed had been partly or fully consumed, or if they presented holes in their surface.

#### Seedling recruitment

Seedling recruitment experiments were set indoors to test the effect of burial depth on recruitment probability. Seeds were buried on 1, 3, 5 and 10 cm and unburied, to test the recruitment probability of seeds that did not go through secondary dispersal. Seeds were watered daily and monitored weekly. We considered recruitment the appearance of the first leaves and ceased monitoring for a given species and depth after two weeks without new recruitments. In total, we used 550 seeds of *P. cattleianum*, 400 of *C. superba* and *N. membranacea* and 180 of *V. bicuhyba*.

### 2.1 Statistical analysis

We estimated the recruitment probability of a seed with diameter *x* dispersed by a primary disperser *j* considering the results from the three experiments. Let *B*_*x,j*_ be the burial depth, *S*_*x,j*_ the seed survival and *R*_*x,j*_ the recruitment. To estimate the final recruitment probability, it is necessary to first determine the burial depth distribution, the seed survival at those different depths and, finally, the recruitment probability of seeds buried at those depths. We set *S*_*x,j*_ = 1 if the seed survived and *S*_*x,j*_ = 0 if it was preyed upon. Similarly, *R*_*x,j*_ = 1 if the seed recruited and *R*_*x,j*_ = 0 otherwise. By taking the conditional probabilities, we derive our equation for the final recruitment probability, represented by *Y*_*i,j*_, as:

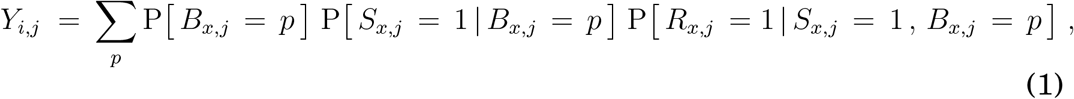

where the sum is carried out over all burial depths *p*. Recall that in our experiment *p* takes the values 0 (unburied), 1, 3, 5 and 10cm.

#### Seed burial

All statistical analysis was done using R 3.6.2 statistical environment (R Development Core Team 2019). To estimate the burial probability we ran a Generalized Linear Model for proportional data (i.e. the proportion of beads that were buried in each station containing 10 piles of feces). For estimating the probability of burial depth, we ran a Cumulative Link Model for ordinal data using the ordinal package (Christensen 2019). As we wanted to calculate the final probability of seedling recruitment considering the results from all experiments (see equation (1)), we categorized the response variable burial depth into the 5 treatment levels from the predation and recruitment experiments: unburied and 1, 3, 5 and 10cm burial depths. In both analysis we did a *χ*^2^ to test the effects of the explanatory variables in the models. We estimated the treatment combinations’ marginal means through the emmeans package (Lenth et al. 2020) as a post-hoc test.

#### Seed predation

We ran several *χ*^2^ tests to evaluate the effect of the explanatory variables: primary seed disperser, seed size and burial depth. For estimating the probability of predation we ran a Generalized Linear Model for proportional data (i.e. the proportion of seeds that were predated in each station containing 10 piles of feces). The differences among treatments were calculated by estimating their marginal means through the emmeans package (Lenth et al. 2020).

#### Seedling recruitment

Finally, to estimate the probability of seed recruitment considering burial depth, we first did a *χ*^2^ test to select the explanatory variables and then ran a Generalized Linear Model for binomial distribution. A post-hoc marginal means test was done to calculate the difference between treatments through the emmeans package (Lenth et al. 2020).

## 3 Results

Seeds with 7mm or more in diameter benefited from howlers’ dispersal (Figure 1). While the proportion of 3mm seeds that recruited varied little with the type of frugivore (18, 20 and 19% for howlers, capuchins and guans, respectively), seeds with 7mm or more had a higher probability of recruiting when dispersed by howlers. The recruitment probability of 7mm seeds was 13% when dispersed by howlers, 8% when dispersed by capuchins and 6% when dispersed by guans. For 10mm seeds, the probability of recruitment was 18% when dispersed by howlers and 8% when dispersed by capuchins or guans. We could not estimate the final seed fate of 14mm seeds because no seed of this class recruited in the recruitment experiment.

**Figure 1:**
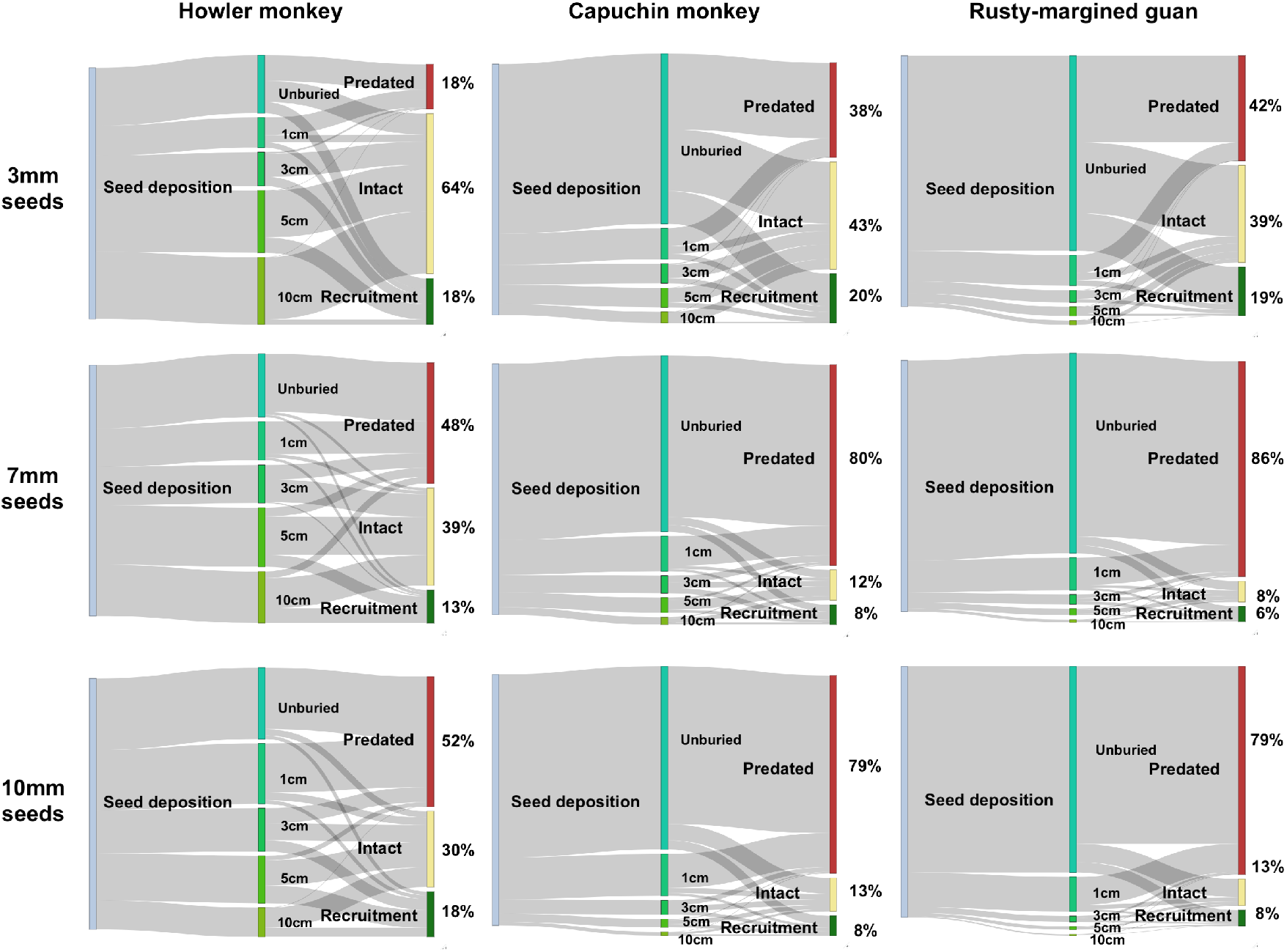
seed fate of 3, 7 and 10mm seeds dispersed by howlers, capuchins and guans. The area in the left of each plot, in grey, represents 100% of deposited seeds. Then, in different shades of green, are the proportions of unburied seeds and seeds buried at 1, 3, 5 or 10cm of depth. Finally, the red, yellow and dark green stripes at the right of the plots are the proportions of seeds with 3, 7 and 10mm of diameter that were preyed upon, remained intact or recruited when dispersed by howlers, capuchins and guans.

The primary disperser and bead size both affected bead burial probability (primary disperser: *χ*^2^ = 317.27, *df* = 2, *p <* 0.001, bead size: *χ*^2^ = 31.14, *df* = 3, *p <* 0.001) and burial depth (primary disperser: *χ*^2^ = 56.51, *df* = 2, *p <* 0.001, bead size: *χ*^2^ = 42.56, *df* = 3, *p <* 0.001). Burial probability was higher for dispersal by howlers in all bead sizes and decreased with increasing bead size (Figure 2). In howler treatments’ bead burial probability varied between 77

**Figure 2:**
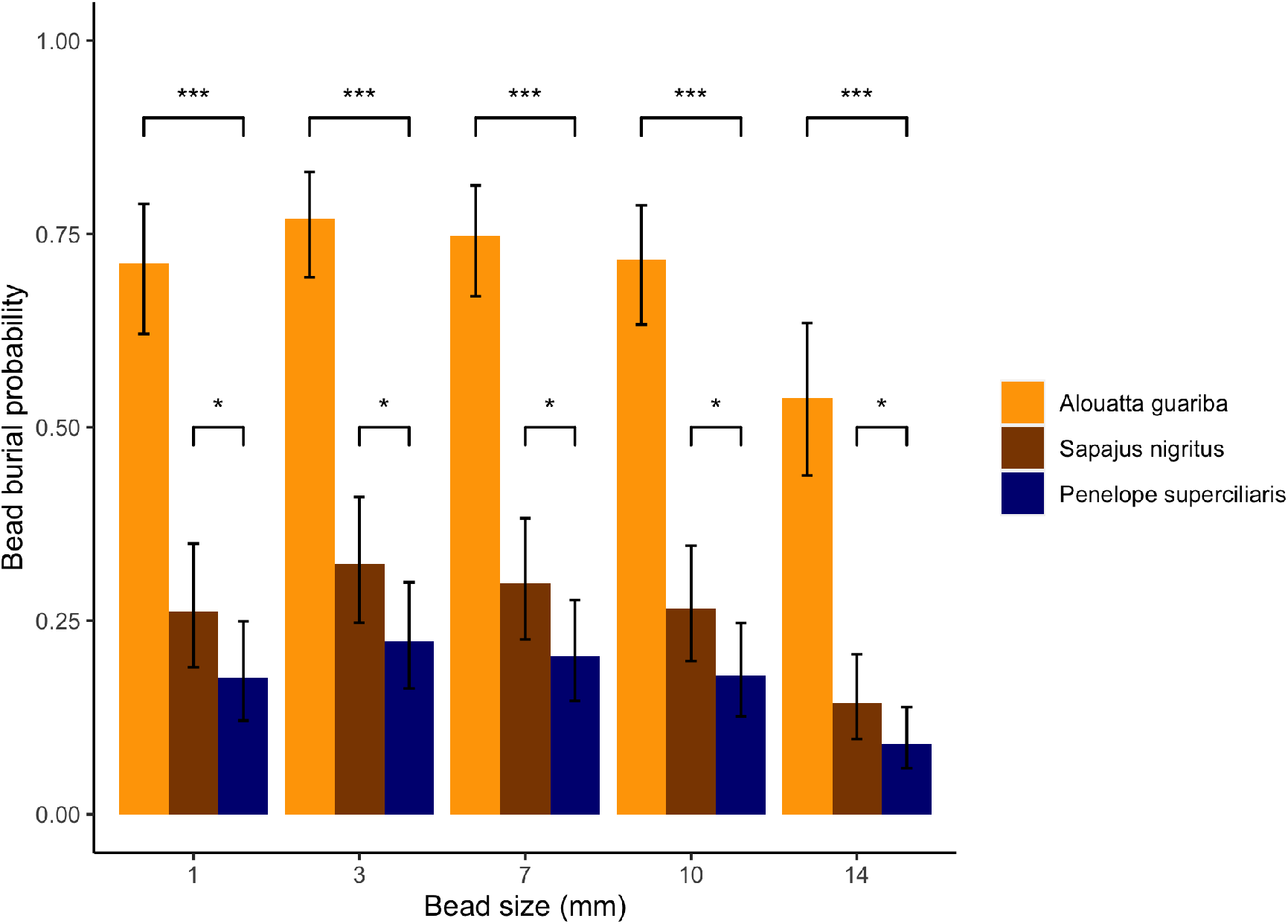
bead burial probability. Percentage of beads buried in howler monkeys’, capuchin monkeys’ and guans’ feces for different bead sizes in orange, brown and blue, respectively. *** corresponds to p < 0.001 and * to p < 0.05.

± 6.8%, for 3mm beads, and 54 ± 10.0%, for 14mm beads. For capuchins’ the probability varied from 32 ± 8.2% to 14 ± 5.4% and for guans’ from 22 ± 6.9% to 9 ± 3.9%, for 3 and 14mm beads, respectively.

Bead burial depth pattern varied according to the primary disperser (Figure 3). In the howlers’ treatments, burial probability increased with depth for 3 (from 16 ± 5.2% to 35 ± 8.5%) and 7mm beads (from 21 ± 6.6% to 27 ± 7.9%), but decreased for 10 (from 34 ± 8,6% to 16 ± 5.6%) and 14mm beads (52 ± 11,9% to 8 ± 4.1%). In contrast, in the capuchins’ and guans’ treatments, burial probability decreased with depth for all bead sizes. When beads were buried in guans’ experiments, it was mostly 1cm deep (from 54.7 ± 5.2%, for 3mm beads, to 88 ± 7.4%, for 14mm beads). In capuchins’ treatments, the difference between probabilities was higher for 3 (from 38 ± 1.0% to 14 ± 5.6%) and 7mm (from 46 ± 12% to 10 ± 4.7%) beads, than for 10 (from 63 ± 11,4% to 5 ± 5.1%) and 14mm beads (from 78 ± 1.0 to 3 ± 1.7%).

**Figure 3:**
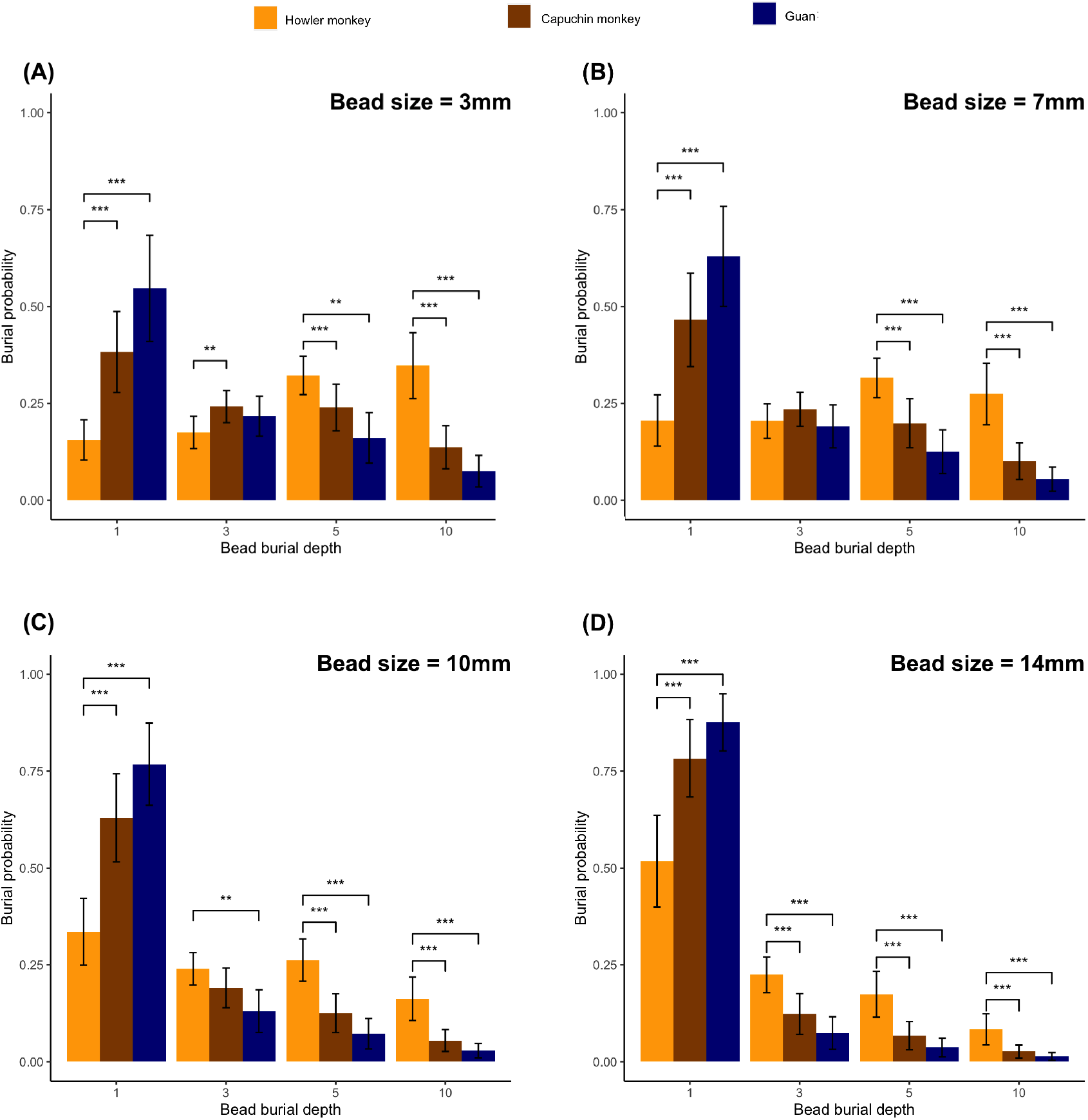
burial depth probability. Percentage of beads buried at 1, 3, 5 and 10cm for different bead sizes. Results for experiments with howler monkeys’, capuchin monkeys’ and guans’ treatments are in orange, brown and blue, respectively. Each bead size is represented in a different plot: (A) 3mm, (B) 7mm, (C) 10mm and (D) 14mm. *** corresponds to p < 0.001, ** to p < 0.005 and * to p < 0.05.

Primary disperser species did not affect predation probability. The model that best predicted predation probability included burial depth (*χ*^2^ = 1519.19, *df* = 4, *p <* 0.001), seed size (*χ*^2^ = 211.26, *df* = 3, *p <* 0.001) and the interaction between these variables (*χ*^2^ = 45.01, *df* = 10, *p <* 0.001). Predation rates decreased with increasing burial depth (Figure 4). However, unburied seeds were not significantly different from seeds buried 1cm deep (*p >* 0.05) in all seed sizes. Predation probability of unburied and 1cm burial varied between 44 ± 6.6% and 58 ± 7.7%, for 3mm seeds, and between 86 ± 4.5% and 81 ± 5.5%, for 7, 10 and 14mm seeds. Almost no predation event occurred for 3mm seeds in depths greater than 1cm. Predation rates did not differ significantly between 7 and 10mm seeds in all burial treatments and it was very rare in seeds buried 5cm and 10 cm deep (lower than 22%).

**Figure 4:**
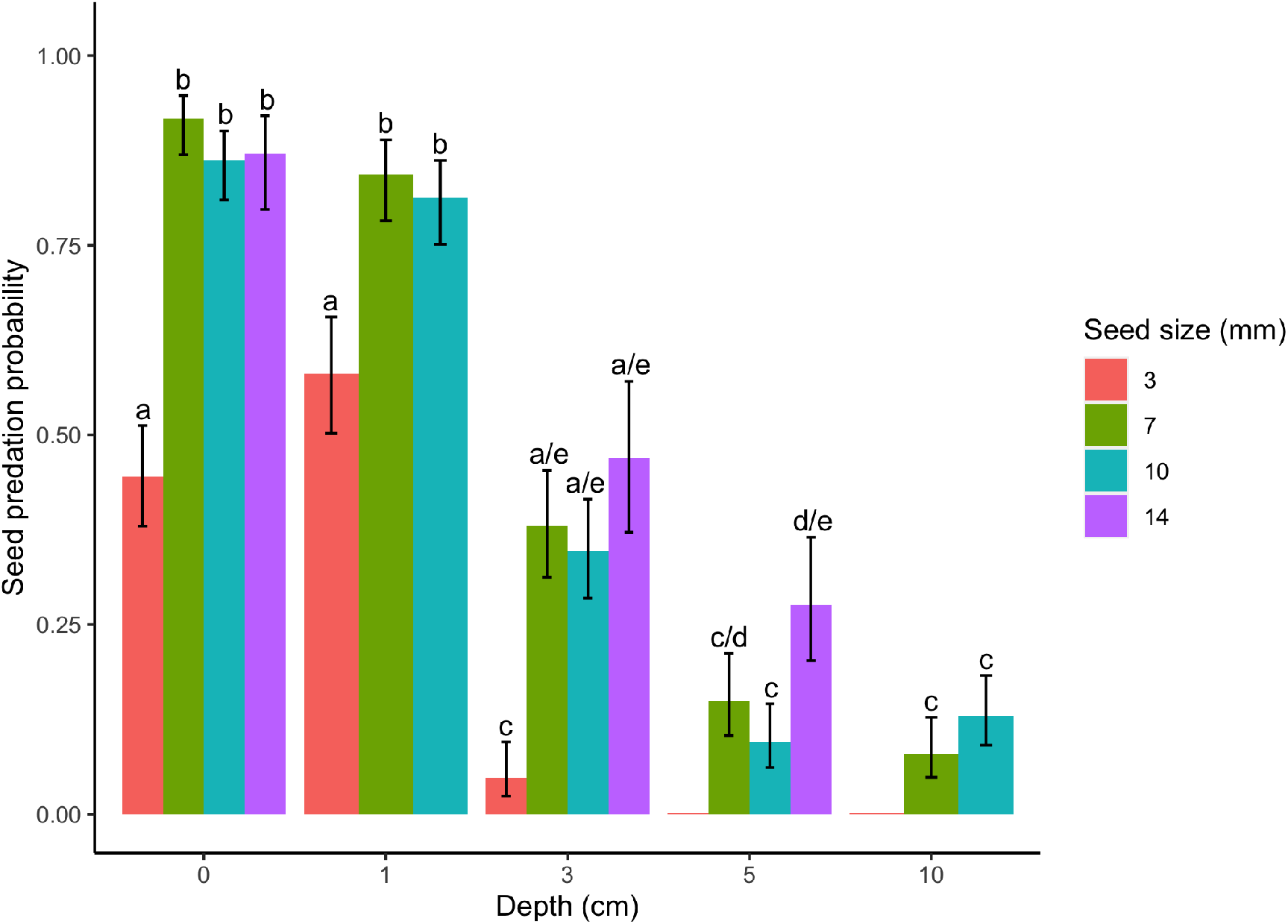
seed predation probability. Predation probability of 3, 7, 10 and 14mm seeds in red, green, blue and purple, respectively. Due to lack of seed quantity, 14mm seeds were not tested for 1 and 10cm depths. 3mm seeds’ predation was tested for 5 and 10cm depths, but no predation event occurred during the period of the experiment.

**Figure 5:**
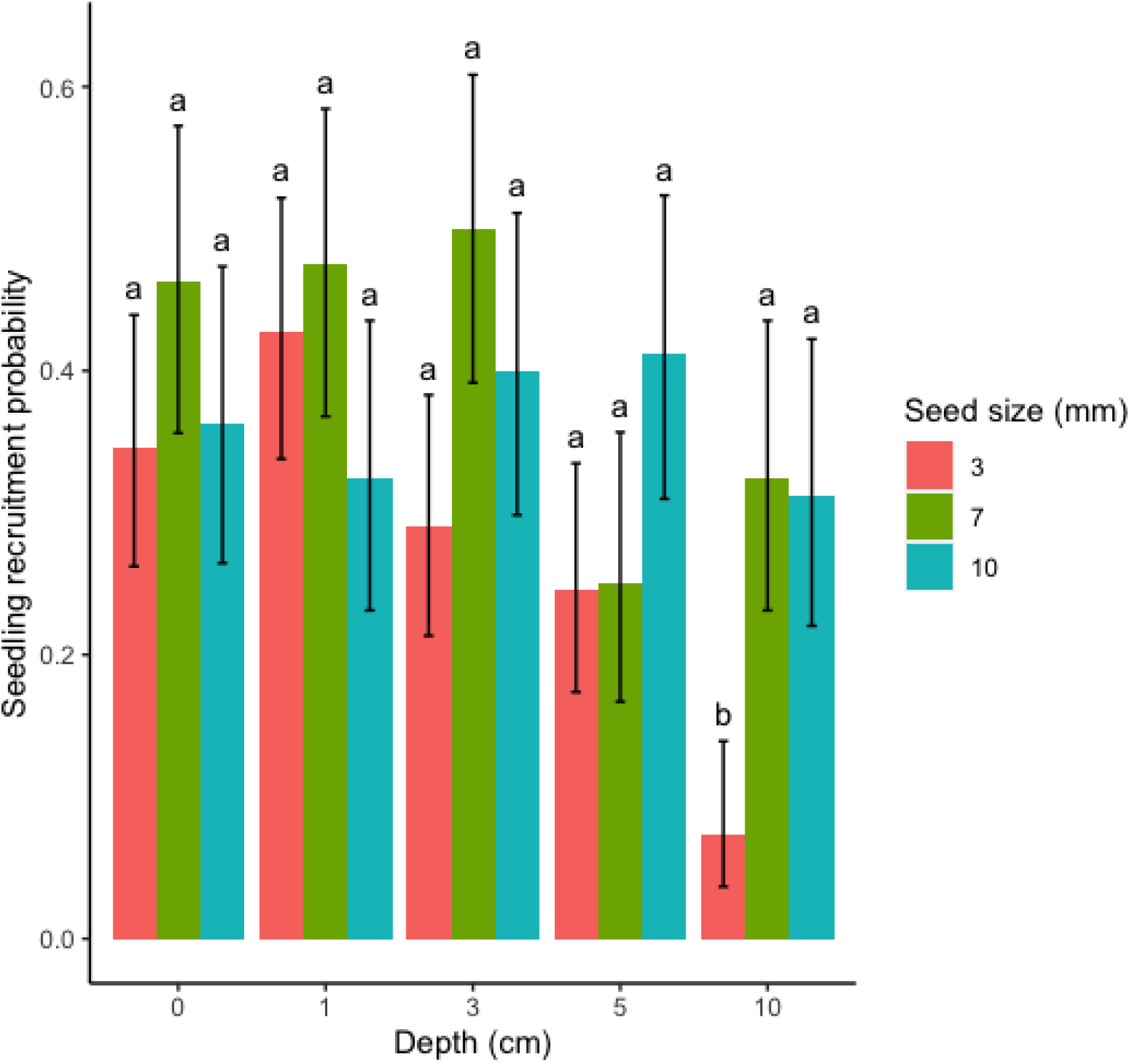
seedling recruitment probability. Recruitment probabilities of 3, 7and 10mm seeds in red, green and blue, respectively. Seeds with 14mm diameter did not recruit at any depth.

As mentioned before, we did not test the effect of primary seed disperser on seedling recruitment. Seed size (*χ*^2^ = 17.95, *df* = 2, *p <* 0.001), burial depth (*χ*^2^ = 32.63, *df* = 4, *p <* 0.001) and the interaction between these variables (*χ*^2^ = 30.24, *df* = 8, *p <* 0.001) affected recruitment probability. Although most treatments did not differ statistically from each other, the effect of burial depth varied with seed size. The recruitment probability of 3mm seeds’ was the one that decreased the most with burial depth. The highest recruitment rate was at 1cm deep (42.7%) and the lowest was at 10cm (7.3%). Seedling recruitment of 7 and 10mm seeds decreased less with burial depth. For seeds with 7mm, the highest recruitment probability was at 3cm deep (50%) and lowest at 5cm deep (25%). For seeds with 10mm, recruitment probability varied little with burial depth, with the highest probability at 5 (41.3%) and lowest at 10cm (31.3%).

## 4 Discussion

Howler monkeys’ seed dispersal in Tijuca National Park improved the recruitment success of large seeds. Seeds dispersed by howler monkeys were buried more often and in greater depths, which increased their survival. However, burial depth tended to decrease the chances of seedling recruitment for all seed sizes. In consequence, the effect of howlers’ seed dispersal differed from capuchins’ and guans’ only for seeds larger than 7mm. Seeds with 3mm had lower predation rates and low recruitment when buried too deep and, thus, did not benefit from howlers’ dispersal. Seeds larger than 7mm, nevertheless, benefited, as their predation rate was higher in shallower depths and recruitment rate didn’t decrease as much when deeply buried.

Dung beetles are assumed to be generalists because they depend on ephemeral and patchily distributed resources. However, recent studies suggest that dung beetles are more attracted to omnivorous mammals’ feces (Whipple and Hoback 2012, Bogoni and Hernandéz 2014, Wurmitzer et al. 2017). In this study, although capuchins are omnivores and howlers have a plant-based diet herbivores, burial probability was significantly higher in howlers’ feces (*circa* 40% higher in all bead sizes). This is probably due to dung beetles being more attracted to large piles of moist dung, similar to the howlers’ defecation pattern, than pelleted depositions, such as the capuchins’ and guans’ (Nichols et al. 2009, Santos-Heredia et al. 2010, Lugon et al. 2017, Raine et al. 2020). Indeed, burial probability in capuchins’ and guans’ feces was more similar (burial probability in capuchins’ feces was *circa* 10% higher than in guans’ feces for all bead sizes). Thus, our results indicate that the amount of dung is a better predictor of attractiveness than dung type, as burial probability was higher in howlers’ feces than capuchins’. Similar results in tropical forests support this idea (Andresen 1999, Culot et al. 2009, 2018).

In accordance with other studies (Andresen 2001, 2002a,b, Santos-Heredia et al. 2010), burial depth was greater in howlers’ treatments, which contained larger piles of dung. Large dung beetles usually are more attracted to larger dung piles due to their active foraging (Peck and Howden 1984). They are more efficient at dung and seed removal (Slade et al. 2007) and bury seeds deeper (Feer 1999, Andresen and Levey 2004). Therefore, one can infer that howler monkey defecations, that are larger and clumped, attract large beetles, which in turn bury seeds deeper.

Burial depth also depended on seed size: smaller seeds were buried deeper in all treatments, as larger seeds are more easily detected and removed by dung beetles. This same result was seen in other studies (Andresen 1999, Feer 1999, Andresen and Levey 2004, Lugon et al. 2017). It should be noted, however, that howlers’ clumped defecation did not attract more seed predators. On the other hand, seed size, burial depth and the interaction between these factors affected predation probability. That is, larger seeds were preyed upon more often and their probability of predation decreased less with increasing depth than for smaller seeds.

Although burial depth decreases seed predation, it was not positively related to seedling recruitment. In this and other studies (reviewed by Andresen and Feer 2005), seedling recruitment had a tendency to decrease with burial depth. However, recruitment probability also depended on seed size: larger seeds recruited more frequently at greater depths. The final outcome was that the proportion of 3mm seeds that recruited did not vary with primary disperser; but, for 7 and 10mm seeds, dispersal by howlers resulted in a higher recruitment success. Even though we had no results for seedling recruitment of 14mm seeds, we can infer that they also benefit from howlers dispersal, as their predation rate at shallow depths was high. Additionally, for all seed sizes, the proportion of seeds that survived but did not recruit was larger when dispersed by howlers. Thus, howlers contribute not only for the recruitment of large seeds but also for the seed bank in TNP.

Ideally, the recruitment experiment should have accounted for primary dispersers. That is, seeds should have been fed to primary dispersers before the experiment, so we could test for the effect of gut passage. We did not include this procedure as we did not have enough seeds. Nonetheless, while howlers have a positive effect on seed germination (Arroyo-Rodríguez et al. 2015), only 34% of seeds remain viable after ingestion by capuchins (Mikich et al. 2015). Thus, the exclusion of gut passage treatment has probably underestimated the positive effect of the reintroduction of howlers in the TNP’s seed dispersal.

The biased extinction of large mammals (Dirzo et al. 2014) and birds (Newbold et al. 2013) is a long-time concern. Larger animals have a disproportionate contribution to seed dispersal (Donoso et al. 2017). For instance, smaller animals that remain in defaunated forests are not able to disperse large seeds. Additionally, as seen in this study, the higher probability of secondary seed dispersal in the piles of dung produced by larger animals can increase the recruitment probability of large seeds. Consequently, the densities of saplings of large-seeded plants are severely reduced in defaunated forests in comparison to small seeded ones (Terborgh et al. 2008). A decreased density of large-seeded plants can have cascading effects on the provisioning of ecosystem services due to the positive relation between carbon storage capacity and seed diameter in tropical trees (Bello et al. 2015).

The loss of large animals also affects dung beetle communities (Bogoni et al. 2019). This trend has been observed specifically for howler monkeys’ presence and abundance (Andresen 2003). Additionally, there is an apparent direction on the reduction of large beetles abundances (Culot et al. 2013). Large beetles have a disproportionate contribution to dung beetle functional activities and a decrease in their abundances can lead to reductions in secondary seed dispersal, bioturbation and nutrient cycling in defaunated areas (Nichols et al. 2009).

Reintroductions can be applied to reverse defaunation scenarios (Oliveira- Santos and Fernandez 2010, Seddon et al. 2014). Yet, most reintroduction efforts are directed towards the conservation of endangered species, and not the restoration of ecological functions and processes (Polak and Saltz 2011). Furthermore, there is a strong bias towards vertebrates reintroductions (Seddon et al. 2005). Reintroductions projects should have a broader view on ecological processes, including all steps and not just the initial interactions. We propose that dung beetles should be reintroduced after the reintroduction of large animals with which they interact. This management action could improve seed dispersal and regeneration of large seeded plants.

Howler monkeys are good candidates for reintroduction projects to restore seed dispersal and plant regeneration. First, they are resilient and able to inhabit and connect small forest patches (Arroyo-Rodríguez and Dias 2010). Secondly, they swallow seeds up to 4.6cm (Arroyo-Rodríguez et al. 2015) in length and although their clumped defecation results in high seed densities, secondary seed dispersal by dung beetles increases large seed survival and recruitment, as discussed above. In fact, seed dispersal by howlers can partially compensate for the absence of larger seed dispersers (Culot et al. 2017). Thus, as howlers deal well even in fragments with a few dozen hectares, they can be the main responsible for the persistence of large-seeded plants in these forest remnants.

The yellow fever outbreak in 2016-2017 (Moreira-Soto et al. 2018) has caused the reduction and extinction of many populations of howlers. The species *Alouatta guariba*, for example, used to be considered of no concern to IUCN and is now vulnerable (Jerusalinsky et al. 2020). Despite howlers’ positive attributes, their reintroduction in TNP was the first with the goal to restore ecological processes (Andresen et al. 2018). Howlers’ reintroduction in TNP was also affected by the yellow fever outbreak: it was interrupted and only 5 individuals are currently in the area. We hope that our results will encourage the resuming of howlers reintroduction in TNP as well as new reintroductions in different areas.

## References

Andresen, E. 1999. Seed Dispersal by Monkeys and the Fate of Dispersed Seeds in a Peruvian Rain Forest. Biotropica 31(1): 145–158.

Andresen, E. 2001. Effects of dung presence, dung amount, and secondary dispersal by dung beetles on the fate of Micropholis guyanensis (Sapotacea) seeds in Central Amazonia. Journal of Tropical Ecology 17: 61–78.

Andresen, E. 2002a. Primary Seed Dispersal by Red Howler Monkeys and the Effect of Defecation Patterns on the Fate of Dispersed Seeds. Biotropica 34(2): 261–272.

Andresen, E. 2002b. Dung beetles in a Central Amazonian rainforest and their ecological role as secondary seed dispersers. Ecological Entomology 27, 257–270.

Andresen, E. 2003. Forest fragmentation on dung beetle communities and functional consequences for plant regeneration. Ecography 26: 87–97.

Andresen, E. and Levey, D. J. 2004. Effects of Dung and Seed Size on Secondary Dispersal, Predation and Seedling Establishment of Rain Forest Trees. Oecologia 139(1): 45–54.

Andresen, E. and Feer, F. 2005. The role of dung beetles as secondary seed disperser and their effect on plant regeneration in tropical rainforests. P.M. Forget, J. E. Lambert, P. E. Hulme and S. B. Vander Wall (eds.): Seed Fate: Predation, Dispersal and Seedling Establishment. Wallingford, UK, CAB International, pp. 331–349.

Andresen, E., Arroyo-Rodríguez, V. and Ramos-Robles, M. 2018. Primate Seed Dispersal: Old and New Challenges. International Journal of Primatology 39: 443–465.

Araujo, B. B. A., Oliveira-Santos, L. G. R., Lima-Ribeiro, M. S., Diniz-Filho, J. A. F. and Fernandez, F. A. S. 2017. Bigger kill than chill: The uneven roles of humans and climate on late Quaternary megafaunal extinctions. Quaternary International 431B, 216–222.

Arroyo-Rodríguez, V. and Dias, P. A. D. 2010. Effects of habitat fragmentation and disturbance on howler monkeys: a review. American Journal of Primatology 72(1): 1–16.

Arroyo-Rodríguez, V., Andresen, E., Bravo, S. P. and Stevenson, P. R. 2015. Seed Dispersal by Howler Monkeys: Current Knowledge, Conservation Implications and Future Directions. M.M. Kowalewski et al. (eds.), Howler Monkeys, Developments in Primatology: Progress and Prospects. New York, Springer Science + Business Media, pp. 111–139.

Bello, C., Galetti, M., Pizo, M. A., Magnago, L. F. S., Rocha, M. F., Lima, R. A. F., Peres, C. A., Ovaskainen, O. and Jordano, P. 2015. Defaunation affects carbon storage in tropical forests. Science Advances 1, e1501105.

Bello, C., Galetti, M., Montan, D., Pizo, M. A., Mariguela, T. C., Culot, L., Bufalo, F., Labecca, F., Pedrosa, F., Constantini, R., Emer, C., Silva, W. R., da Silva, F. R., Ovaskainen, O. and Jordano, P. 2017. Atlantic frugivory: a plant-frugivore interaction data set for the Atlantic Forest. Ecology 98, 1.

Bogoni, J. A. and Hernandéz, M. I. M. 2014. Attractiveness of Native Mam-mals’ Feces of Different Trophic Guilds to Dung Beetles. Journal of Insect Science 14(299): 1–7.

Bogoni, J. A., Giovâni, P. and Peres, C. A. 2019. Co-declining mammal-dung beetle faunas throughout the Atlating Forest biome of South America. Ecography 42: 1803–1818.

Bufalo, F. S., Galetti, M. and Culot, L. 2016. Seed Dispersal by Primates and Implications for the Conservation of a Biodiversity Hotspot, the Atlantic Forest of South America. Int J Primatol 37: 333–349.

Chapman, C. A. and Dunham, A. E. 2018. Primate Seed Dispersal and Forest Restoration: An African Perspective for a Brighter Future. Int J Primatol 39, 427–442.

Christensen, R. H. B. 2019. ordinal—Regression Models for Ordinal Data. R package version 2019.12-10. https://CRAN.R-project.org/package= ordinal.

Cid, B., Figueira L., Mello, A., Pires, A. S. and Fernandez, F. A. S. 2014. Shortterm success in the reintroduction of the red-humped agouti Dasyprocta leporina, an important seed disperser, in a Brazilian Atlantic Forest reserve. Tropical Conservation Science 7:796–810.

Culot, L., Huynen, M.C., Gérard, P. and Heymann, E. W. 2009. Short-term post-dispersal fate of seeds defecated by two small primate species (Saguinus mystax and Saguinus fuscicollis) in the Amazonian forest of Peru. Journal of Tropical Ecology 25: 229–238.

Culot, L., Bovy, E., Vaz-de-Mello, F. Z., Guevara, R. and Galetti, M. 2013. Selective defaunation affects dung beetle communities in continuous Atlantic rainforest. Biological Conservation 163: 79–89.

Culot, L., Bello, C., Batista, J. L. F., do Couto, H. T.Z. and Galetti, M. 2017. Synergistic effects of seed disperser and predator loss on recruitment success and long-term consequences for carbon stocks in tropical rainforests. Scientific Reports 7: 7662.

Culot, L., Huynen, M. C. and Heymann, E. 2018. Primates and Dung Beetles: Two Dispersers Are Better than One in Secondary Forest. International Journal of Primatology 39, 397–414.

Dehling, D. M., Jordano, P., Schaefer, H. M., Böhning-Gaese and Schleuning, M. 2016. Morphology predicts species’ functional roles and their degree of specialization in plant-frugivore interactions. Proc. R. Soc. B. 283: 20152444.

Dirzo, R., Young, H. S., Galetti, M., Ceballos, G., Isaac, N. J. B. and Collen, B. 2014. Defaunation in the Anthropocene. Science 345(6195): 401–406.

Donoso, I., Schleuning, M., García, D., and Fründ, J. 2017. Defaunation effects on plant recruitment depend on size matching and size trade-offs in seed-dispersal networks. Proc. R. Soc. B 284: 20162664.

Eisenberg J. F. and Thorington R. W. 1973. A preliminary analysis of a neotropical mammal fauna. Biotropica 5:150–161.

Estrada, A., Garber, P. A., Rylands, A. B., Roos, C., Fernandez-Duque, E., Di Fiore, A., Nekaris, K. A. I., Nijman, V., Heymann, E. W., Lambert, J. E., Rovero, F., Barelli, C., Setchell, J. M., Gillespie, T. R., Mittermeier, R. A., Arregotia, L. V., de Guinea, M., Gouveia, S., Dobrovolski, R., Shanee, S., Shanee, N. Boyle, S.A., Fuentes, A., MacKinnon, K. C., Amato, K. R., Meyer, A. L. S., Wich, S. Sussman R. W., Pan, R., Kone, I. and Li, B. 2017. Impending extinction crisis of the world’s primates: Why primates matter. Science Advances 3: e1600946.

Feer, F. 1999. Effects of dung beetles (Scarabaeidae) on seeds dispersed by howler monkeys (Alouatta seniculus) in the French Guianan rainforest. Journal of Tropical Ecology 15, 129–142.

Freitas, S. R., Neves, C. L. and Chernicharo, P. 2006. Tijuca National Park: two pioneering restorationist initiatives in Atlantic Forest in southeastern Brazil. Braz J Biolo 66(4): 975–982.

Fuzessy, L. F., Cornelissen, T. G., Janson, C. and Silveira, F. A. O. 2016. How do primates affect seed germination? A meta-analysis of gut passage effects on neotropical plants. Oikos 125: 1069–1080.

Fuzessy, L. F., Janson, C. H. and Silveira, F. A. O. 2017. How far do Neotropical primates disperse seeds? Am J Primatol 79:e22659.

Genes, L., Cid, B., Fernandez, F. A. S. and Pires, A. S. 2017. Credit of ecological interactions: A new conceptual framework to support conservation in a defaunated world. Ecology and Evolution 7: 1891–2897.

Genes, L., Fernandez, F. A. S., Vaz-de-Mello, F. Z., da Rosa, P., Fernandez, E. and Pires, A. S. 2019. Effects of howler monkey reintroduction on ecological interactions and processes. Conservation biology 33(1): 88–89.

ICMBio (Instituto Chico Mendes de Conserva/ccaõ da Biodiversidade. 2008. Plano de Manejo: Parque Nacional da Tijuca. Instituto Brasileiro de Desenvolvimento Florestal, Brasília, Brasil.

Jerusalinsky, L., Cortes-Ortíz, L., de Melo, F. R., Miranda, J., Alonso, A. C., Buss, G., Alves, S. L., Bicca-Marques, J., Neves, L., Ingberman, B., Fries, B., da Cunha, R., Mittermeir, R. A. and Talebi, M. 2020. Alouatta guariba. The IUCN Red List of Threatened Species 2020: e.T39916A17926390.

Julliot, C. 1996. Seed dispersal by red howler monkeys (Alouatta seniculus) in the tropical rain forest of French Guiana. International Journal of Primatology 17: 239–258.

Koch, P. L. and Barnosky, A. D. 2006. Late Quaternary Extinctions: State of the Debate. Annu. Rev. Ecol. Evol. Syst. 37:215–50.

Lenth, R. V., Buerkner, P., Herve, M., Love, J., Riebl, H. and Singmann, H. 2020. Estimated Marginal Means, aka Least-Squares Means. R package version 1.5.3. https://github.com/rvlenth/emmeans/issues.

Lugon, A. P., Boutefeu, M., Bovy, E., Vaz-de-Mello, F. Z., Huynen, M. C., Galetti, M. and Culot, L. 2017. Persistence of the effect of frugivore identity on post-dispersal seed fate: consequences for the assessment of functional redundancy. Biotropica 49(3): 293–302.

McConkey, K. R. and Brockelman, W. Y. 2011. Nonredundancy in the dispersal network of a generalist tropical forest tree. Ecology 92(7): 1492–1502.

Mikich, S. B. 2002. A dieta frugívora de Penelope superciliaris (Cracidade) em remanescnetes de floresta estacional semidecidual no centro-oeste do Paraná, Brasil e sua relaçaõ com Euterpe edulis (Arecaceae). Ararajuba 10(2):207–217.

Mikich, S. B., Liebsch, D., Almeida, A. and Miyazaki, R. D. 2015. O papel do macaco-prego Sapajus nigritus na dispersaõ de sementes e no controle potencial de insetos-praga em cultivos agrícolas e florestais. In L. M. Parron, J. R. Garcia, E. B. de Oliveira, G. G. Brown and R.B. Prado (eds.): Serviços Ambientais em Sistemas Agrícolas e Florestais do Bioma Mata Atlântica. Brasília, DF, Embrapa, pp. 257–265.

Mittelman, P., Kreischer, C., Pires, A. S. and Fernandez, F. A. S. 2020. Agouti reintroduction recovers seed dispersal of a large-seeded tropical tree. Biotropica 00, 1–9.

Moreira-Soto, A., Torres, M. C., de Mendonça, M. C. L., Mares-Guia, M. A. M., Rodrigues, C. D. S., Fabri, A. A., dos Santos, C.C., Araújo, E.S.M., Fischer, C., Nogueira, R. M. R., Drosten, C., Sequeira, P. C., Drexler, J. F. and Filippis, A. M. B. 2018. Evidence for multiple sylvatic transmission cycles during the 2016-2017 yellow fever virus outbreak, Brazil. Clinical Microbiology and Infection 24(9): 1019.e1-2019.e4.

Newbold, T., Scharlemann, J., Butchart, S., Şekercioğlu, Ç., Alkemade, R., Booth, H. and Purves, D. 2013. Ecological traits affect the response of tropical forest bird species to land-use intensity. Proc. R. Soc. B 280, 20122131.

Nichols, E., Gardner, T. A., Peres, C. A., Spector, S. and The Scarabaeinae Research Network. 2009. Co-declining mammals and dung beetles: an impending ecological cascade. Oikos 118: 481–487.

Oliveira-Santos, L. G. R and Fernandez, F. A. S. 2010. Pleistocene rewilding, frankenstein ecosystems, and an alternative conservation agenda. Conservation Biology 24, 2–5.

Padua, J. A. 2002. Um sopro de destruiçaõ: pensamento político e crítica ambiental no Brasil escravista. 1786-1888. Zahar, Rio de Janeiro, Brazil.

Pardini, R., Bueno, A. d. A., Gardner, T. A., Prado, P. I. and Metzger, J. P. 2010. Beyond the fragmentation threshold hypothesis: regime shifts in biodiversity across fragmented landscapes. Plos One 5.

Peck, S. B. and Howden, H. F. 1984. Response of a Dung Beetle Guild to Different Sizes of Dung Bait in a Panamanian Rainforest. Biotropica 16(3): 235–238.

Peres, C. 2002. Synergistic Effects of Subsistence Hunting and Habitat Fragmentation on Amazonian Forest Vertebrates. Conservation Biology 15(6): 1490–1505.

Peres, C. and Palacios, E. 2007. Basin-Wide Effects of Game Harvest on Vertebrate Population Densities in Amazonian Forests: Implications for Animal-Mediated Seed Dispersal. Biotropica 39(3): 304–315.

Polak, T. and Saltz, D. 2011. Reintroduction As an Ecosystem Restoration Technique. Conservation Biology 25(3): 424–427.

Development Core Team, R. 2019. R: A language and environment for statistical computing. Vienna: R Foundation for Statistical Computing http://www.R-project.org/.

Raine, E. H. and Slade, E. M. 2019. Dung beetle-mammal associations: methods, research trends and future directions. Proceedings B 286: 20182002.

Raine, E. H., Mikich, S. B., Lewis, O. T and Slade, E. M. 2020. Linking dung beetle-mediated functions to interactions in the Atlantic Forest: Sampling design matters. Biotropica, 52: 215–220.

Ripple, W. J., Wolf, C., Newsome, T. M., Betts, M. G., Ceballos, G., Courchamp, F., Hayward, M. W., Van Valkenburgh, B., Wallach, A. D. and Worm, B. 2019. Are we eating the world’s megafauna to extinction? Conservation Letters 12:e12627.

Santos-Heredia, C., Andresen, E. and Zárate, D. A. 2010. Secondary seed dispersal by dung beetles in a Colombian rain forest: effects of dung type and defecation pattern on seed fate. Journal of Tropical Ecology 26(4): 355–364.

Schleuning, M., Fründ, J. and García, D. 2015. Predicting ecosystem functions from biodiversity and mutualistic networks: an extension of trait-based concepts to plant-animal interactions. Ecography 38:001–013.

Scholtz, C. H., A. L. V. Davis, and U. Kryger, editors. 2009. Evolutionary Biology and Conservation of Dung Beetles. Pensoft, Bulgaria.

Seddon, P. J., Soorae, P. S. and Launay, F. 2005. Taxonomic bias towards reintroduction projects. Animal Conservation 8, 51–58.

Seddon, P. J., Griffiths, C. J., Soorae, P. S. and Armstrong, D. P. 2014. Reversing defaunation: Restoring species in a changing world. Science 345(6195): 406–412.

Shepherd, V. E. and Chapman, C. A. Dung beetles as secondary seed dispersers: impact on seed predation and germination. Journal of Tropical Ecology 14, 199–215.

Slade, E. M., Mann, D. J., Villanueva, J. F. and Lewis, O. T. 2007. Experimental Evidence for the Effects of Dung Beetle Functional Group Richness and Composition on Ecosystem Function in a Tropical Forest. Journal of Animal Ecology 76, 1094–1104.

Svenning, J. -C., Pedersen, P. B. M., Donlan, C. J., Ejrnæs, R., Faurby, S., Galetti, M., Hansen, D. M., Sandel, B., Sandom, C. J., Terborgh, J. W. and Vera, F. W. M. 2016. Science for a wilder Anthropocene: synthesis and future directions for trophic rewilding research. PNAS 113, 898–906.

Terborgh, J., Nunez-Iturri, G., Pitman, N. C. A., Valverde, F. H. C., Alvarez, P.,Swamy, V.. Pringle, E.G. and Paine, C. E. T. 2008. Tree recruitment in an empty forest. Ecology 89(6): 1757–1768.

Traveset, A. and Riera, N. 2005. Disruption of a Plant-Lizard Seed Dispersal System and Its Ecological Effects on a Threatened Endemic Plant in the Balearic Islands. Conservation Biology 19(2): 421–431.

Vidal, M. M., Pires, M. M. and Guimarães Jr., P.R. 2013. Large vertebrates as the missing components of seed-dispersal networks. Biological Conservation 163: 42–48.

Wang, B. C. and Smith, T. B. 2002. Closing the seed dispersal loop. Trends in Ecology and Evolution 17(8): 379–386.

Whipple, S.D. and Hoback, W. W. 2012. A Comparison of Dung Beetle (Coleoptera: Scarabaeidae) Attraction to Native and Exotic Mammal Dung. Environ. Entomol. 41(2): 238–24.

Wurmitzer, C., Blüthgen, N. Krell, F.T., Maldonado, B., Ocampo, F., Müller, J.K. and Schmitt, T. 2017. Attraction of dung beetles to herbivore dung and synthetic compounds in a comparative field study. Chemoecology 27: 75–84.

